# Transcriptome analysis of IPF fibroblastic foci identifies key pathways involved in fibrogenesis

**DOI:** 10.1101/2020.03.10.984955

**Authors:** Delphine Guillotin, Adam R. Taylor, Manuela Platé, Paul F. Mercer, Lindsay M. Edwards, Ross Haggart, Gino Miele, Robin J. McAnulty, Toby M. Maher, Robert E. Hynds, Mariam Jamal-Hanjani, Richard P. Marshall, Andrew J. Fisher, Andy D. Blanchard, Rachel C. Chambers

**Author notes:** indicates that these first authors contributed equally to this work (Delphine Guillotin, Adam R. Taylor, Manuela Platé). indicates that these senior authors contributed equally to this work (Andy D. Blanchard, Rachel C. Chambers).

## Abstract

Fibroblastic foci (FF) represent the cardinal pathogenic lesion in idiopathic pulmonary fibrosis (IPF) and comprise activated fibroblasts and myofibroblasts, the key effector cells responsible for dysregulated extracellular matrix deposition in multiple fibrotic conditions. The aim of this study was to define the major transcriptional programmes involved in fibrogenesis in IPF by profiling un-manipulated myo/fibroblasts within FF *in situ* by laser capture microdissection.

The challenges associated with deriving gene calls from low amounts of RNA and the absence of a meaningful comparator cell type were overcome by adopting novel data mining strategies and by using weighted gene co-expression network analysis (WGCNA), as well as an *eigengene*-based approach to identify transcriptional signatures which correlate with fibrillar collagen gene expression. WGCNA identified prominent clusters of genes associated with cell cycle, inflammation/differentiation, translation and cytoskeleton/cell adhesion. Collagen *eigengene* analysis revealed that TGF-β1, RhoA kinase and the TSC2/RHEB axis formed major signalling clusters associated with collagen gene expression. Functional studies using CRISPR-Cas9 gene edited cells demonstrated a key role for the TSC2/RHEB axis in regulating TGF-β1-induced mTORC1 activation and collagen I deposition in mesenchymal cells reflecting IPF and other disease settings, including cancer-associated fibroblasts. These data provide strong support for the human tissue-based and bioinformatics approaches adopted to identify critical transcriptional nodes associated with the key pathogenic cell responsible for fibrogenesis *in situ* and further identifies the TSC2/RHEB axis as a potential novel target for interfering with excessive matrix deposition in IPF and other fibrotic conditions.

**What is the key question?:** Can we identify a transcriptional signature associated with collagen gene expression in the fibrotic focus, the cardinal fibrotic lesion in IPF?

**What is the bottom line?:** We herein define the major transcriptional programmes involved in fibrogenesis in IPF by profiling myo/fibroblasts within FF *in situ* by laser capture microdissection.

**Why read on?:** The data provide strong support for a human tissue-based approach to identify critical transcriptional nodes associated with fibrogenesis *in situ* and further identifies the TSC2/RHEB axis as a potential novel target for interfering with excessive matrix deposition in IPF and other fibrotic conditions.

## INTRODUCTION

Fibrosis, defined as the abnormal accumulation of extracellular matrix (ECM), is a pathological feature of several major chronic inflammatory and metabolic diseases, including idiopathic pulmonary fibrosis (IPF), the most rapidly progressive and fatal fibrotic conditions with a median survival of 3.5 years from diagnosis.[1] Fibrosis can also influence cancer progression as part of the stromal response to the tumour.[2] Current anti-fibrotic drugs, pirfenidone and nintedanib, approved for the treatment of IPF, slow rather than halt disease progression and have significant side-effect profiles.[3, 4] There therefore remains an urgent need to develop novel therapeutic strategies for IPF and other fibrotic conditions.

The aetiology of IPF remains poorly defined but the current favoured hypothesis proposes that IPF arises as a result of a highly dysregulated wound healing response following repeated epithelial injury in response to an environmental inciting agent in genetically susceptible and aged individuals. IPF has a distinct histopathological pattern of usual interstitial pneumonia (UIP) in which fibroblastic foci (FF), comprising aggregates of activated fibroblasts and myofibroblasts embedded in a collagen-rich extracellular matrix, represent discrete sites of lung injury, repair and active fibrogenesis.[5] These lesions represent a key histological diagnostic feature of UIP and their abundance is related to disease progression in IPF.[6] In order to shed novel light on the mechanisms leading to pathogenic fibrogenesis in IPF, we sought to define the transcriptional signature of these cardinal fibrotic lesions.[6] While single cell RNA sequencing (scRNAseq) has been reported in IPF,[7] this approach does not discriminate between profiles obtained from cells within FF or from other non-involved locations within the lung. Laser capture microdissection (LCM) of IPF lung tissue, provides an ideal approach to capture and profile unperturbed myo/fibroblasts exclusively from within FF. Comparator cells are challenging to define given the potential diverse origins and phenotypes of myo/fibroblasts in IPF and importantly the paucity of interstitial fibroblasts which can be captured from healthy control lung. Moreover, recent scRNASeq data available on bioRχiv,[8] suggest that “pathological ACTA2-expressing IPF myofibroblasts represent an extreme pole of a continuum connected to a quiescent ACTA2-negative stromal population rather than resident or activated fibroblasts in control lung”. We herein report on the generation of a composite gene expression profile from more than 60 individual FF derived from 13 patients. We adopt a novel informatics approach which obviates the need for a comparator group, based on the principles of weighted gene coexpression network analysis (WCGNA). We further employ a collagen *eigengene* approach[9] to define clusters of genes which functionally correlate with collagen I and III gene expression and therefore inform on the critical transcriptional programme underpinning pathogenic fibrogenesis. Key signalling nodes identified within the collagen *eigengene* cluster comprised gene signatures associated with the master regulator of fibrogenesis, transforming growth factor β1 (TGF-β1), as well as the TSC2/RHEB axis. This axis is a well characterised molecular switch for the activation of mechanistic target of rapamycin complex 1 (mTORC1), a key molecular node that integrates metabolic, energy, hormonal, and nutritional signals to regulate downstream cellular responses, including proliferation, growth, metabolism and protein synthesis.[10] Functional studies using CRISPR-Cas9 gene-edited primary human lung fibroblasts (pHLFs) confirmed the involvement of RHEB in TGF-β1-mediated mTORC1 activation and collagen I deposition. This observation was generalizable to other mesenchymal cells, including hepatic stellate cells (HSCs) and cancer associated fibroblasts (CAFs). Taken together the findings reported provide strong support for the transcriptomic approach adopted to identify functionally relevant pathways involved in pathogenic fibrogenesis *in situ*. Our data further suggest that the TSC2/RHEB axis might represent a potential therapeutic target for interfering with fibrogenesis in the setting of fibrosis and the stromal reaction in cancer.

## METHODS

### Patient material

Frozen IPF lung tissue was obtained from patients either undergoing lung transplantation for end–stage disease (n=10) or surgical lung biopsy for diagnostic purposes (n=3). Patients with IPF were diagnosed in accordance with current international guidelines.[11] The human biological samples were sourced ethically and their research use was in accord with the terms of the informed consents (11/NE/0291; 12/EM/0058; 10/H0504/9).

### Laser capture microdissection and microarray analysis

Frozen OCT embedded samples were cryo-sectioned into 8μm sections and processed with a modified haematoxylin and eosin (H&E) stain.[12] FF were identified by immunostaining a serial guide slide for αSMA and captured using a PALM MicroBeam 4 Laser Microdissection microscope (Zeiss). Care was taken to avoid capturing overlying epithelium (visualized by immunostaining for CK7 on a second serial guide slide). 3-6 captures (~0.015 mm^2^) per sample were collected. RNA was extracted using the Picopure RNA Isolation kit (Life Technologies) and processed for hybridization to Affymetrix HG-U133_plus_2.0 microarrays.

### RNAscope^®^ *in situ* hybridization

All RNAscope^®^ analyses were performed on independent IPF lung tissue (Advanced Cell Diagnostics) using a standard RNAscope^®^ 2.5 HD Red protocol according to the manufacturer’s instructions.

### WGCNA and collagen *eigengene* analysis

WGCNA was performed on the 9035 probe sets defined as the composite IPF FF signature in R v3.3.3.[9] Briefly, we selected a power (β) of 4 (the minimum power with R^2^ > 0.9) to construct the co-expression network from which the topological overlap matrix was calculated. To generate gene modules we used *minModuleSize* = 200 and default cutHeight which yielded 16 modules. Modules with *eigengene* correlations R >|0.7| were merged using the *mergeCloseModules* function leaving 14 independent modules. Biological function to the modules was assigned using MetaCore™ (Thomson Reuter) pathway enrichment. Sankey plots were generated to describe the biological function of modules.

### Statistical Analysis

Statistical analysis of gene expression data was performed in R v3.3.3. Correlation coefficients were calculated using the Pearson method (*base*). WGCNA was performed using the wgcna package. P-Values for correlations were calculated using *corPvalueStudent* function in the *wgcna* package. Enrichment analysis and network construction were performed in R v3.3.3 using the *metabaseR* package (Thompson Reuters™) database version 6.36.69400. The Girvan-Newman algorithm[13] was used to identify subnetworks in the collagen module.

### *In vitro* experiments

Control and IPF pHLFs (REC reference 12/EM/0058) and CAFs from lung adenocarcinoma (via the Tracking Cancer Evolution through Therapy (TRACERx) clinical study, REC reference 13/LO/1546) were grown from explant cultures as previously described.[14] Primary adult HSCs were obtained from Zen-Bio (#HP-F-S). RNAseq was performed on control pHLFs as described in [15] (GSE102674). CRISPR-Cas9 gene editing followed the protocol described in [14] using a guide RNA sequence targeting *RHEB* (AGATGCCGCAGTCCAAGTCCCGG). Protein phosphorylation and expression were assessed by western blotting as described in [14] using p-4EBP1 (#13443), p-P70S6K (#9234), 4EBP1 (#9644), P70S6K (#9202) and RHEB (#13879) antibodies (all from Cell Signalling). α-tubulin (#9099) was used as a loading control. Type I collagen deposition under macromolecular crowding conditions was assessed by high-content imaging as described in [16]. Procollagen production was based on levels of hydroxyproline quantified by reverse phase high-performance liquid chromatography (HPLC) as described in [17]. I*n vitro* data were analysed by two-way ANOVA with Tukey multiple-comparisons testing (Graphpad Prism). Data were considered statistically significant at p⍰<⍰0.05.

## RESULTS

### Identification of an *in situ* gene expression signature for IPF fibroblastic foci

In order to generate an *in situ* gene expression signature for IPF FF, lesions from 13 flash-frozen IPF tissue samples (Table 1; Figure 1a) were captured by LCM (please see Methods section). RNA was isolated from 61 captures, amplified and cDNA hybridized to Affymetrix microarrays. To interrogate the cellular purity of the captures we examined the normalized microarray intensity values of myofibroblast (*ACTA2*) and epithelial (*CDH1, CK7, EpCAM*) marker genes (Figure 1b). The large difference in median Log2 intensity values obtained for myofibroblast and epithelial cell markers provided assurance regarding the quality of the captures. We next validated and examined the spatial localization of selected matrisomal mRNAs detected within the FF transcriptome in additional patient samples by selecting genes representing a range of microarray expression intensities (*SPARC, ELN, POSTN*) using RNAScope^®^ *in situ* hybridization. An RNAScope^®^ signal was detectable for all three genes within FF but not in the overlying epithelium (Figure 1c). Finally, we took advantage of a recently available scRNASeq dataset submitted on bioRχiv to interrogate enrichment for IPF fibroblast and myofibroblast marker genes.[8] The fibroblast marker genes, HAS1 and HAS2, were detected in less than 7 patients and were therefore excluded from our FF signature. The myofibroblast markers genes, COL8A1 and ACTA2, were detected in 13/13 and in 12/13 patients, respectively. These data suggest that myofibroblasts are enriched in IPF FF.

**Table 1.**
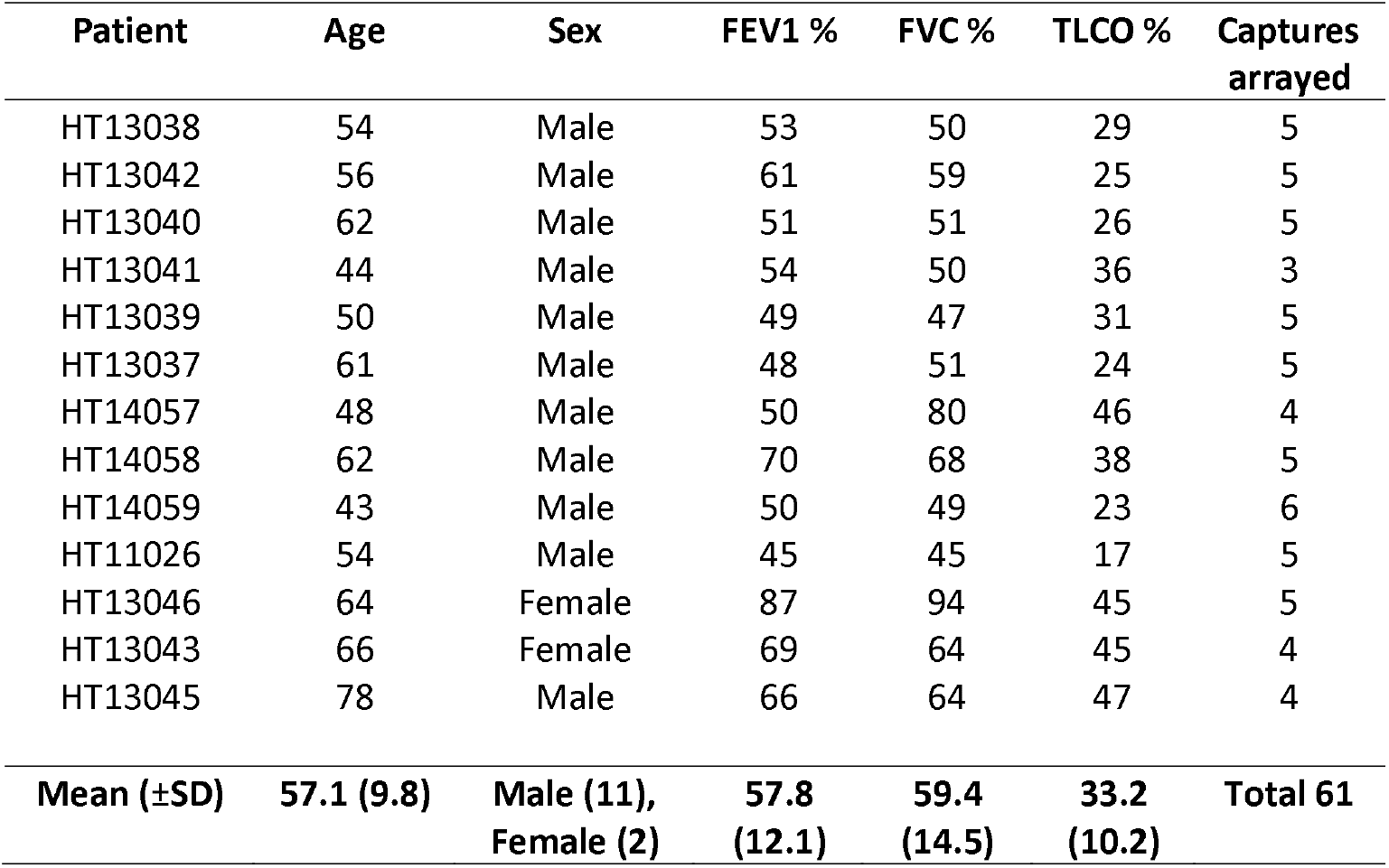
Baseline characteristics of IPF patients for tissues used for laser capture microdissection. The demographics of the patients included in this study are shown. All patients were diagnosed with IPF in line with ATS/ERS 2011 guidelines. Pulmonary function tests were measured pre-transplantation or pre-biopsy and expressed as percentage of the predicted value. Data are shown as mean (±) SD. Definition of abbreviations: FEV1 = forced expiratory volume in one second; FVC = forced vital capacity; TLCO = transfer factor of the lung for carbon monoxide.

**Figure 1.**
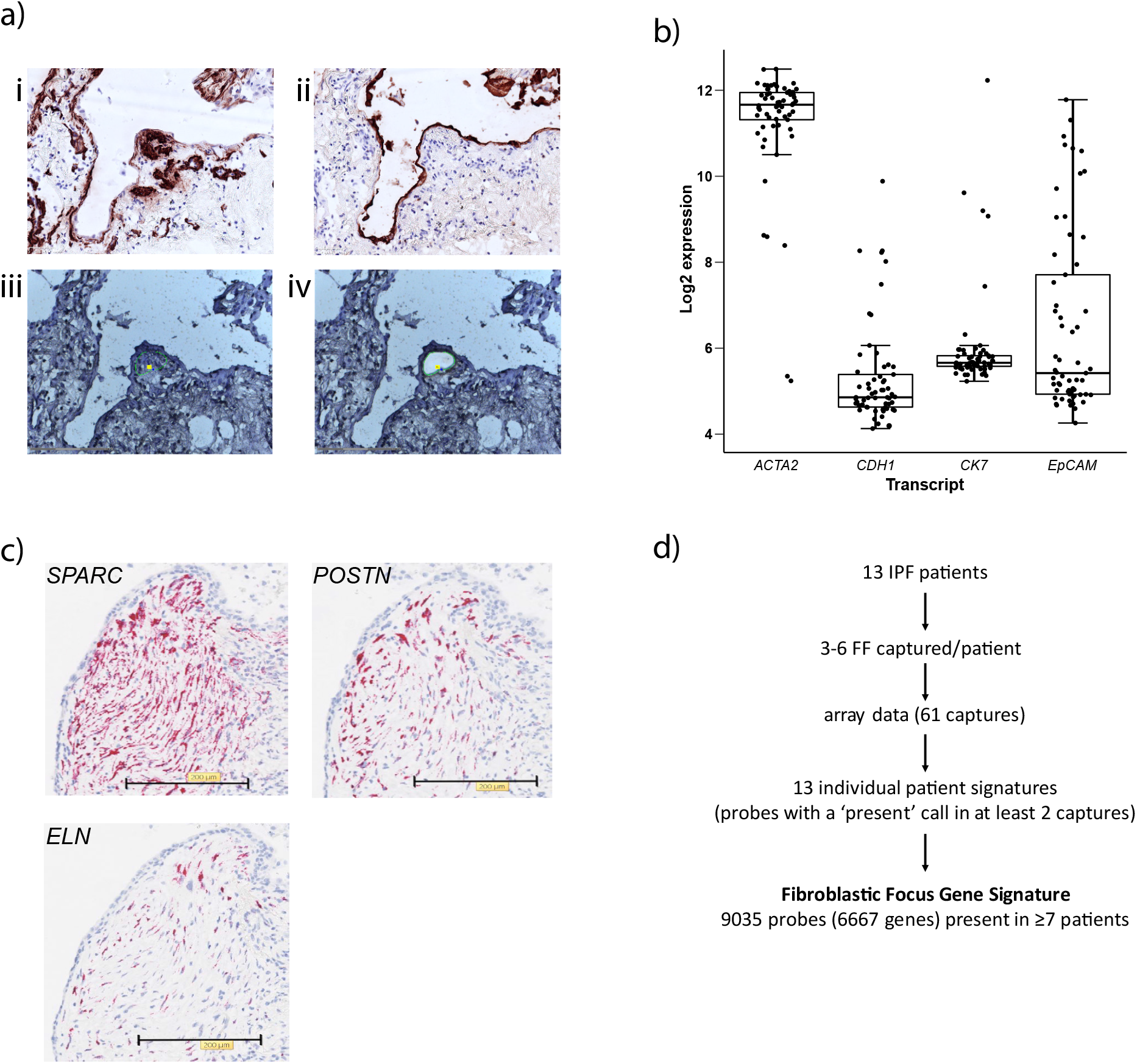
Transcriptomic profiling of laser captured fibroblastic foci from IPF patients. **a)** Captures were obtained from fresh frozen IPF lung tissue obtained from 13 patients. αSMA-rich FF were identified by αSMA immunostaining and captured from serially stained H&E sections, taking care to avoid the overlying CK7-positive epithelium. Representative sections of αSMA (i) and CK7 (ii) staining are shown, as well as sections before (iii) and after (iv) laser capture of a typical fibroblastic focus. **b)** Median Log2 intensity values for myofibroblast (*ACTA2*) and epithelial (*CDH1, CK7, EpCAM*) marker genes. **c)** Confirmation of spatial expression of selected genes within FF in independent patient samples by RNAScope^®^ *in situ* hybridization (SPARC, POSTN, ELN). **d)** Generation of the fibroblastic focus gene expression signature. For each of the 13 patient lung samples, fibroblastic foci were captured, the RNA was extracted, amplified and cDNA was arrayed on an Affymetrix HG-U133_plus_2.0microarray. Using the presence call information on the array, we excluded probes that were present in less than two captures for the same patient (approximately 5% error rate). The final probeset intensity was averaged per patient. Probesets detected in the majority (≥7) of the patient signatures were included to generate a final IPF FF gene expression signature/transcriptome.

To generate an *in situ* IPF FF transcriptome, we first generated independent gene signatures for each of the 13 IPF patients using an informatic pooling strategy (described in Figure 1d). We then derived a composite IPF FF transcriptomic signature by selecting probesets expressed in the majority (≥7) of individual IPF patient signatures. This cut-off identified 9035 probesets corresponding to 6667 individual genes. The complete dataset is available on Gene Expression Omnibus (GSE98925) (table S1).

### Weighted gene co-expression network analysis (WGCNA) of the *in situ* FF transcriptome

In order to identify transcriptional programmes and biological pathways active in myo/fibroblasts within FF, we applied WGCNA and Metacore™ analyses of the composite FF transcriptome. WGCNA identified 14 independent co-expression modules reflecting sets of highly correlated gene transcripts (Figure 2a, Table S1). We next aimed to identify modules with the highest proportion of genes upregulated in IPF compared with control lungs by cross-referencing published whole lung microarray expression data (GSE10667).[18] The WGCNA turquoise module comprised genes with the most significant and highest average fold-change in IPF versus control (Figure 2b-c) and was therefore considered to represent the most enriched module with potential disease-relevant biology in the laser-captured FF.

**Figure 2:**
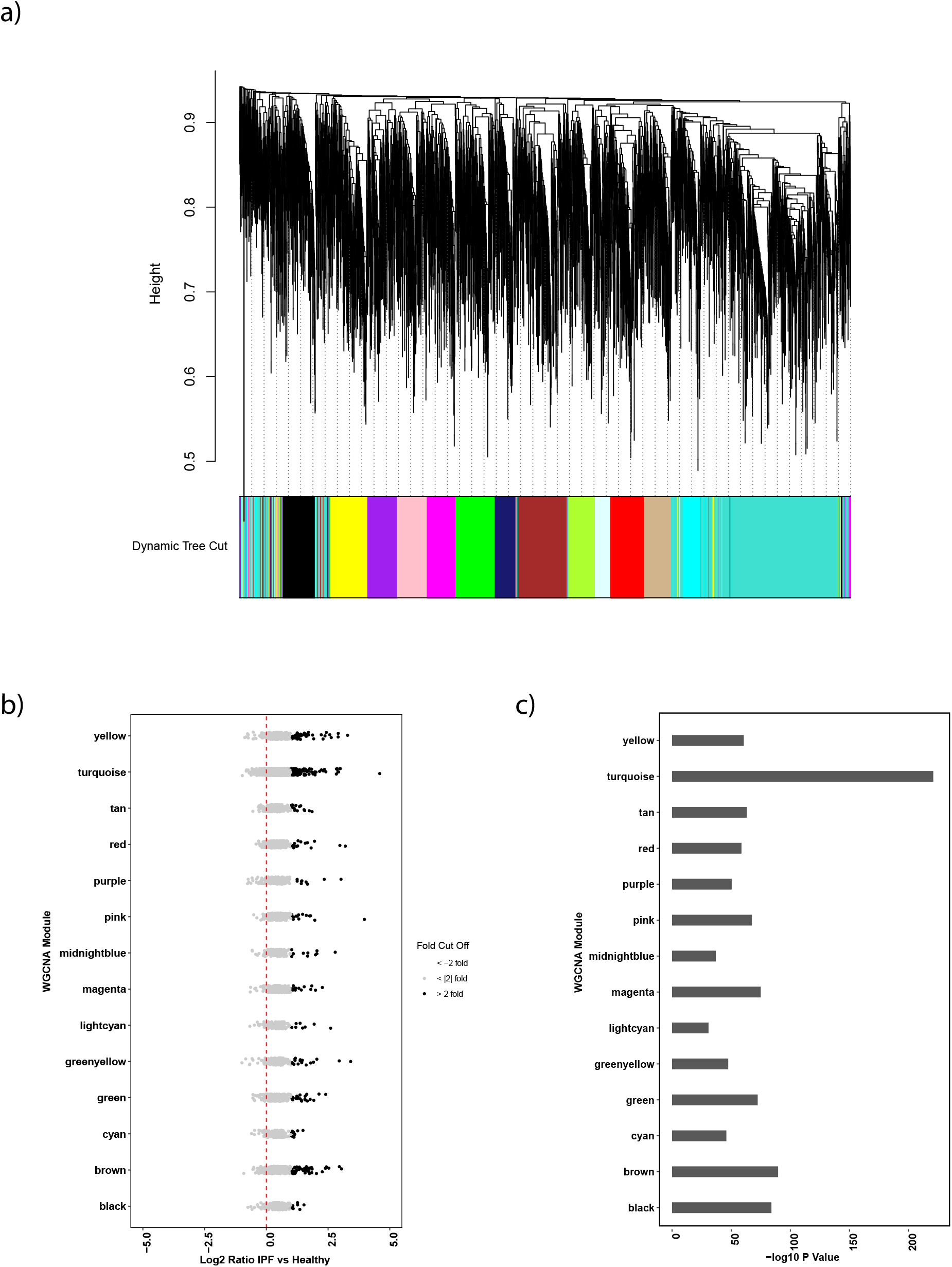
Weighted Gene Co-Expression Analysis (WGCNA) of the composite fibroblastic focus transcriptome. **a)** The composite FF transcriptome was subjected to WGCNA and the gene dendrogram is shown identifying 14 independent co-expression modules. Modules can appear in multiple regions of the dendrogram as highly correlated, smaller modules are collapsed. **b)** The gene expression in IPF versus healthy lung from published data (GSE10667) for all genes of each module was interrogated. The log2 ([diseased]/[healthy]) ratios are shown with > 2-fold difference shown in black. **c)** −log10 p-values from a two-sided t-test performed on each module log2 ratios, to determine which sets were significantly different from zero.

Within the 2607 genes represented in the WGCNA turquoise module, we mapped 2051 Metacore™ network objects (representing genes, proteins, complexes and metabolites) and constructed a single biological network comprising 1021 IPF-enriched genes. In order to deconvolute this network further, we identified sub-clusters (i.e. local neighbourhoods) and investigated the seven local neighbourhoods containing more than 20 genes (Figure 3a). Pathway analysis of these local neighbourhoods identified key genes that were most influential in the pathway mapping and may therefore represent important regulatory nodes in FF (Figure 3b-c). This included genes representing known cardinal features of the activated myofibroblast, such as cytoskeletal remodelling (*RHOA, RAC1, ROCK, ARHGEF1*), as well as highlighting JAK/STAT and NFκB signalling pathways and a small number of transcription factors.

**Figure 3:**
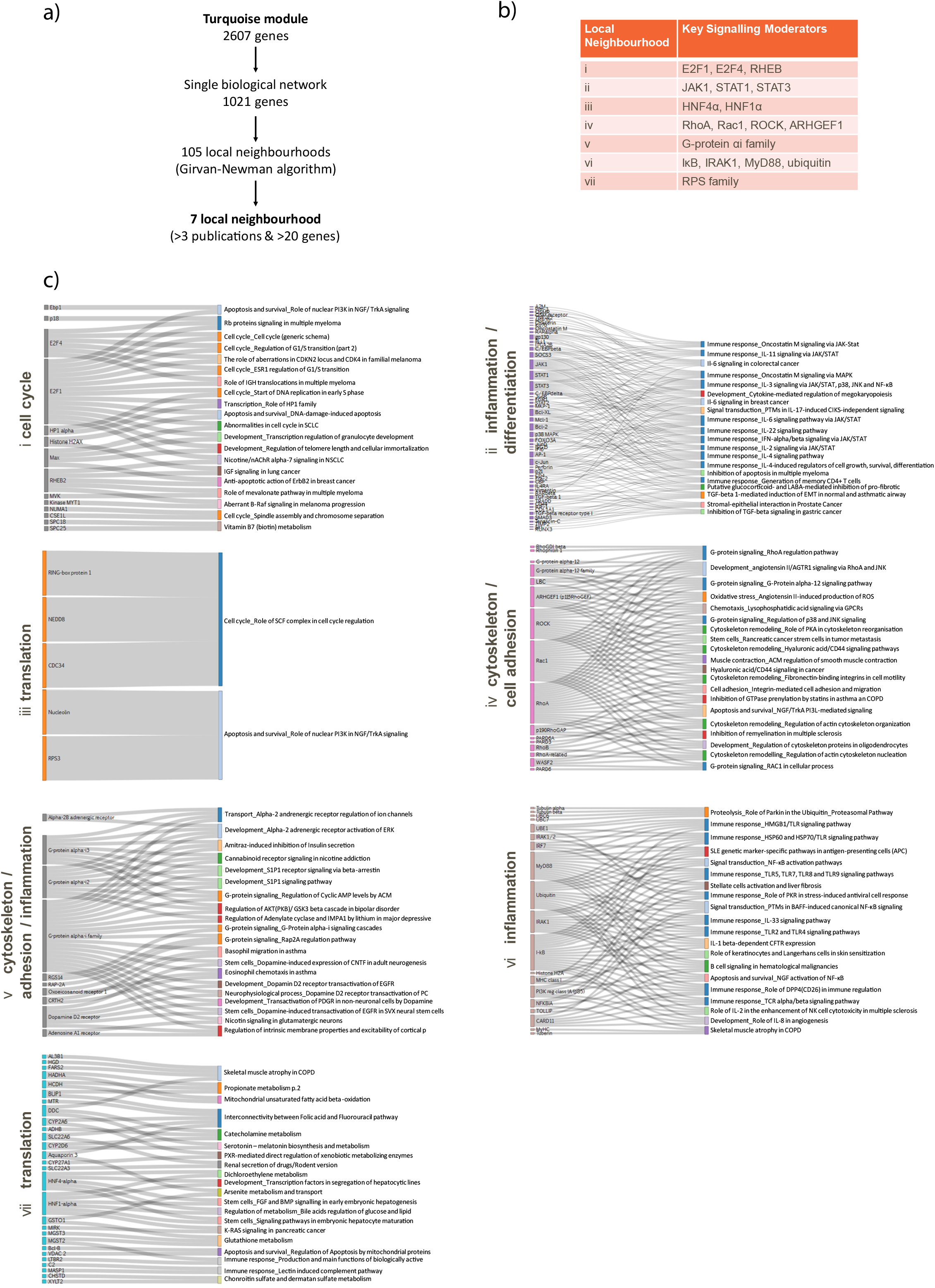
Network and pathway analysis of the WGCNA turquoise module. **a)** A biological network was constructed from the 2607 genes in the turquoise WGCNA module. After removal of the edges supported by less than three citations, 7 local neighbourhoods containing at least 20 genes were investigated. **b)** Pathway analysis of the seven sub-clusters is shown displaying the genes that appear most frequently in pathways identified to represent key modulators of neighbourhood biology. **c)** The pathways detected with the highest significance are shown in detail as Sankey diagrams.

### Collagen *eigengene* analysis of the *in situ* IPF FF transcriptome identifies key transcriptional programmes associated with fibrogenesis

We next sought to define the transcriptional programmes and therefore the key pathways associated with pathogenic collagen production in IPF. To this end, we adapted the core principle of WGCNA to create a bespoke collagen *eigengene* from the first principal component of *COL1A* and *COL3A* probesets and then correlated the rest of the microarray probesets making up the FF signature. 362 probesets, representing 300 individual genes, correlated either positively or negatively with the collagen *eigengene* (Figure 4a, Table S1). Fifteen genes correlated with an R>|0.9|, including four genes that were highly *inversely* correlated with collagen gene expression.

**Figure 4:**
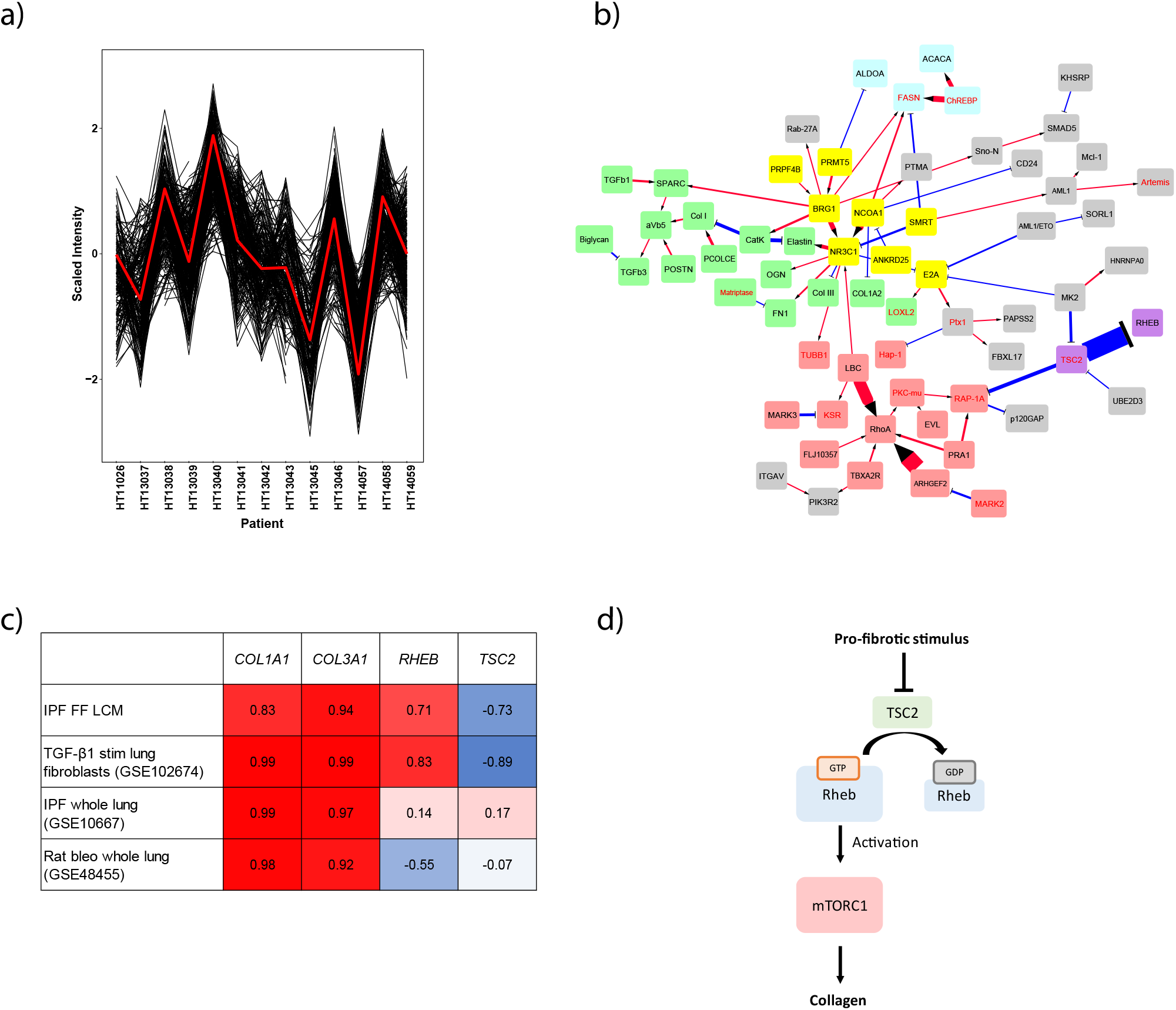
Collagen *eigengene* analysis. **a)** The expression of *COL1A* and *COL3A* mRNA probes across the 13 patient profiles was used to create a collagen *eigengene* (red line) and all other probesets correlating to this (p<0.01), assigned as collagen correlating genes (black lines). Array expression data is shown as Z-transformed log2 intensity. Z-transformed log2 intensities of negatively correlated genes were inverted for illustration. b) A single interaction network was derived from the 300 genes forming the collagen *eigengene* as represented by three or more citations and excluding implied interactions. The network includes a TGF-β1/ECM module (green), a Rho kinase module (red), a transcriptional module (yellow), a metabolism module (blue) and a TSC/RHEB module (purple). The proteins are linked with a red arrow to represent activation or a blue line to represent an inhibition with the thickness of the lines reflecting the existing body of evidence for the interaction. Gene names illustrated in black font were positively correlated with the collagen *eigengene*, whereas genes in red were negatively correlated. c) Table reflects correlation coefficients for *COL1A1*, *COL3A1*, *TSC2* and *RHEB* with comparable *eigengenes* (*COL1A1* and *COL3A1*) calculated independently in the captured IPF FF (IPF FF LCM) and in previously published datasets for TGF-β1 stimulated human primary lung fibroblasts, IPF whole lung and in the rat bleomycin model. **d)** Figure illustrates a proposed model for mTORC1 activation in response to a pro-fibrotic stimulus (e.g. TGF-β1) or a fibrotic microenvironment (e.g. FF). TSC2 (−ve) and *RHEB* (+ve) expression correlations relative the collagen *eigengene* would favour persistent mTORC1 activation and collagen deposition via GTP-bound Rheb.

To further characterize this collagen *eigengene* module, we constructed an interaction network using MetaCore™ which we refined to a network of 66 genes following removal of predicted rather than experimentally-validated interactions and those without directionality (Figure 4b). This network featured genes that represent TGF-β1/ECM biology (green) and RhoA associated cytoskeletal remodelling (red), supporting this network as a core regulatory feature of the FF myo/fibroblast phenotype. Additional modules included transcriptional regulation (yellow) and cell metabolism (blue), as well as a small module featuring *TSC2* and *RHEB*, a well-characterised molecular switch for the activation of mTORC1 signalling. RHEB, which encodes the GTPase and proximal activator of mTORC1, *RHEB*, correlated positively with the collagen *eigengene* (R +0.71). In contrast, *TSC2* which encodes tuberin within the TSC complex, a major allosteric inhibitor of RHEB, was inversely correlated (R −0.73). We next determined whether a similar TSC2/RHEB motif was represented in vitro in our published RNAseq dataset of TGF-β1-stimulated fibroblasts (GSE102674) and in published whole lung IPF and rat bleomycin model gene expression datasets (GSE10667 and GSE48455).[15, 18] A comparable *eigengene* was calculated using *COLA1* and *COLA3* expression levels and correlation coefficients for *COL1A1*, *COL3A1*, *TSC2* and *RHEB* were generated. As expected *COLA1* and *COLA3* expression correlated across all datasets; whereas the specific *RHEB* and *TSC2* correlations were only present in TGF-β1-stimulated fibroblasts but not in datasets from whole fibrotic human and murine lung, where transcriptional profiles represent a composite derived from multiple cell types (Figure 4c). The LCM FF and TGF-β1-stimulated fibroblast correlations would therefore support a potential model favouring mTORC1 activation (Figure 4d).

### Functional validation of the TSC2/RHEB node during fibrogenesis *in vitro*

Given the robust correlation of the TSC2/RHEB node with the collagen *eigengene* in IPF FF and in TGF-β1 activated myofibroblasts *in vitro*, we next sought to determine whether TSC2/RHEB represents a potential molecular switch involved in regulating collagen synthesis in TGF-β1-stimulated pHLFs. To this end, we generated *RHEB*-deficient pHLFs using CRISPR-Cas9 gene editing. Monitoring the phosphorylation of the mTORC1 downstream substrates, P70S6K and 4E-BP1 (Serine 65) at 3 hours following TGF-β1 stimulation in control and *RHEB*-deficient fibroblasts revealed that RHEB was critical for TGF-β1-induced mTORC1 activation (Figure 5a). Analysis of procollagen synthesis, based on the quantitation of hydroxyproline, further demonstrated that RHEB was necessary for TGF-β1-induced procollagen synthesis (Figure 5b). This observation was validated by specifically assessing collagen I deposition in response to TGF-β1 stimulation under macromolecular crowding conditions by high-content imaging (Figure 5c, d). Finally, we also measured procollagen synthesis following TGF-β1 stimulation in IPF pHLFs, HSCs and CAFs. In each case, there was an almost complete inhibition of TGF-β1-induced collagen production in *RHEB*-edited cells (Figure 6).

**Figure 5:**
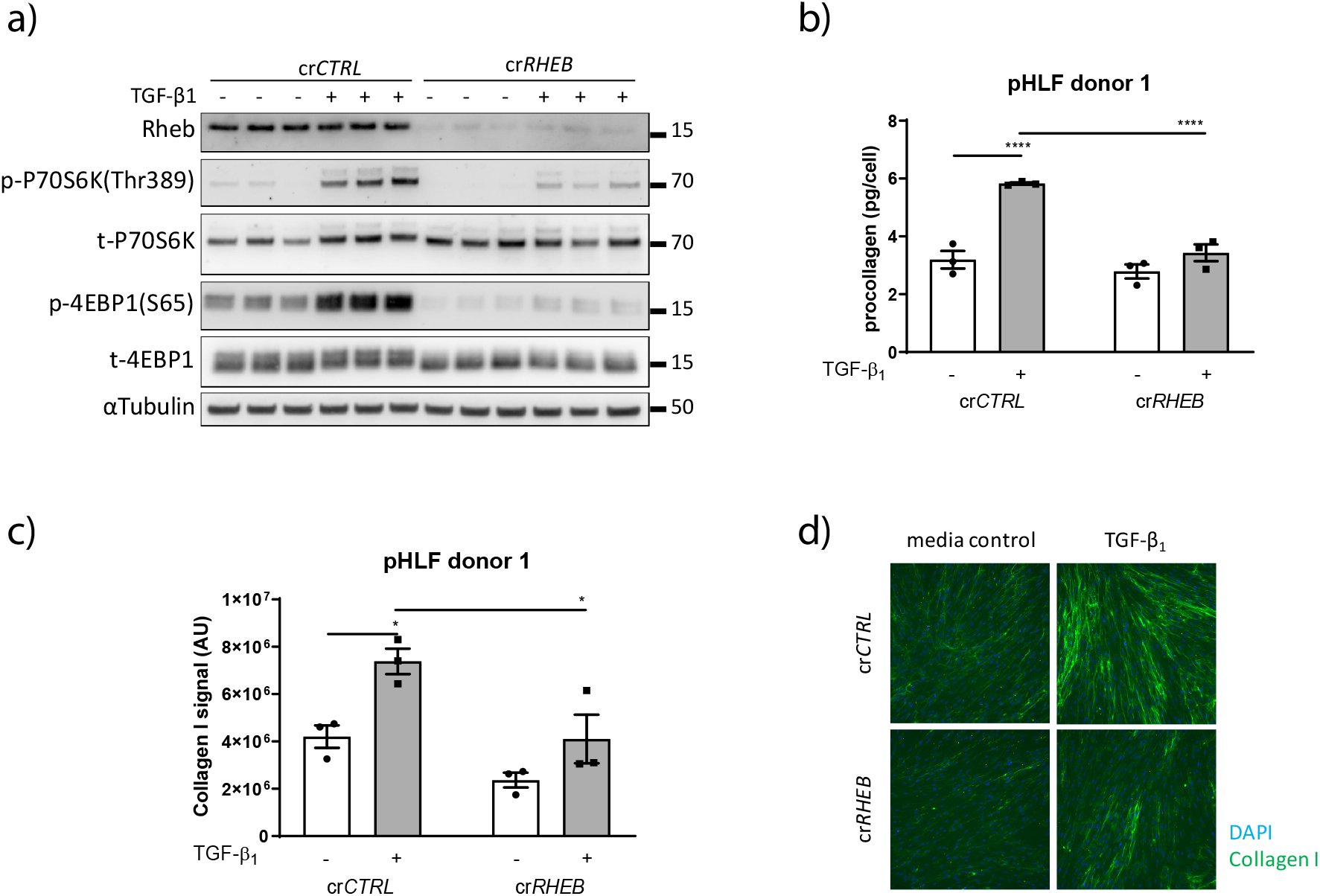
RHEB is essential for TGF-β1-stimulated mTORC1 activation and collagen production in primary human lung fibroblasts (pHLFs). **a)** CRISPR-Cas9 gene editing was used to disrupt RHEB in pHLFs. A predesigned control gRNA sequence was used to generate matched wild-type pHLFs (cr*CTRL*). RHEB-disrupted (crRHEB) and cr*CTRL* pHLFs were stimulated with TGF-β1 (1ng/mL) or media alone for 3 hrs. Successful gene editing of *RHEB* was confirmed by immunoblotting. mTORC1 signalling was evaluated by monitoring the phosphorylation of the mTORC1 downstream substrates, P70S6K and 4E-BP1 by immunoblotting. **b)** Procollagen production was measured by quantifying hydroxyproline levels in ethanol-insoluble proteins in cell supernatants after 48 hrs stimulation with TGF-β1 (1ng/mL) by reverse-phase HPLC and normalised to the cell count. **c)** Collagen I deposition after 48hrs stimulation with TGF-β1 (1ng/mL) was measured by high-content imaging of collagen I immunofluorescence in cells grown under macromolecular crowding conditions. Data are represented as collagen I signal normalised to cell count (n=4 fields per well). **d)** Representative images of collagen I deposition are shown. Data in graphs are presented as mean (±) SEM of 3 replicate wells per condition. Differences between groups were evaluated by two-way ANOVA with Tukey multiple comparison testing, * = p<0.05, **=p<0.01, ***=p<0.001, ****=p<0.0001.

**Figure 6:**
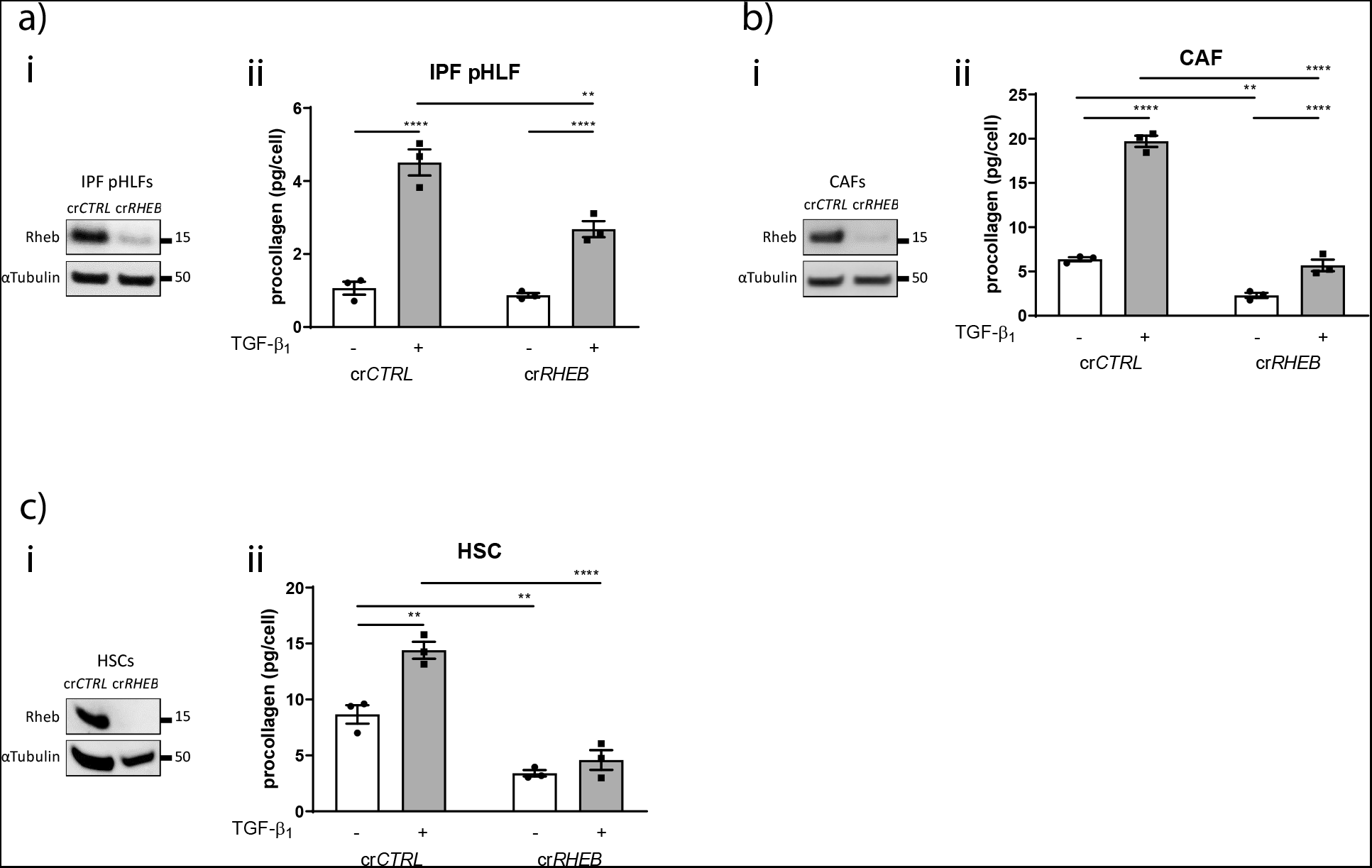
RHEB is essential for TGF-β1-stimulated collagen deposition in IPF pHLFs, CAFs and HSCs. CRISPR-Cas9 gene editing was used to disrupt RHEB (cr*RHEB*) and a cr*CTRL* was used as a control in: **a)** IPF pHLF; **b)** primary CAFs derived from lung adenocarcinoma; and **c)** primary HSCs. (i) Successful RHEB gene editing was confirmed by assessing RHEB levels by immunoblotting and (ii) procollagen production was measured by quantifying hydroxyproline levels in ethanol-insoluble proteins in cell supernatants after 48 hrs following stimulation with TGF-β1 (1ng/mL) by reverse-phase HPLC and normalised to the cell count. Data are presented as mean (±) SEM of 3 replicate wells per condition. Differences between groups were evaluated with two-way ANOVA with Tukey multiple comparison testing, * = p<0.05, **=p<0.01, ***=p<0.001, ****=p<0.0001.

## DISCUSSION

The aim of this study was to identify key transcriptional pathways driving fibrogenesis in IPF by fingerprinting myo/fibroblasts located within FF, the defining pathological lesion of IPF/UIP. The use of informatics-based analysis of gene expression data to identify genes, pathways and mechanisms of relevance in organs and tissues of interest has become an established path to uncovering important pathobiology.[19] scRNAseq methodologies are rapidly advancing and recent studies have provided important insights into the heterogeneity and diversity of mesenchymal, epithelial and alveolar macrophage populations in the setting of pulmonary fibrosis.[20–22] However, spatial information, and therefore the precise anatomical location from which the cells are derived, is lost during single cell dissociation. It is also worth commenting that the cell dissociation process can also potentially influence gene expression profiles.[21] In order to generate an unperturbed gene expression signature for IPF myo/fibroblasts and identify pathways associated with pathological collagen expression within the cardinal lesion in IPF,[6] we chose to adopt an LCM-based approach which was optimised to overcome the challenges associated with isolating high quality RNA from a small number of cells embedded within a collagen-rich fibrotic lesion. This allowed us to identify a a unique set of 6667 genes representing a generic transcriptome of the FF.

Our bioinformatics approach obviated the need for a comparator cell type, in light of recent findings indicating that pathological ACTA2-expressing IPF myofibroblasts represent an extreme pole of a continuum connected to a quiescent ACTA2-negative stromal population rather than resident or activated fibroblasts in control lung. It is not possible to capture sufficient number of myo/fibroblasts from the interstitium of uninjured normal adult lung. The lack of a suitable comparator cell type to identify differentially expressed genes in FF was obviated by using contemporary bioinformatic approaches. WGCNA has become widely adopted to identify groups of genes with similar expression profiles (modules) in large high-dimensional genomic datasets. The functional correlation of genes aligned by expression within an *eigengene* underlies the principle of the WGCNA methodology where individual modules represent discrete functions.[9] These modules highlight related biological mechanisms and can provide insights into disease pathomechanisms. The largest WCGNA turquoise module identified within our dataset comprised 2607 genes and was the most significantly enriched module in terms of genes previously identified to be upregulated in whole IPF lungs compared to healthy controls.[18] RhoA/Rac1/ROCK signalling was prominent within the WGCNA turquoise module and this feature is consistent with the coordinated cytoskeletal rearrangements associated with highly contractile cells, such as myofibroblasts.[23] Strong representation of NFκB signalling in FF was not as predictable but adds to previous evidence implicating this signalling hub in the context of liver fibrosis[24, 25] and further provides support for a role for FF myo/fibroblasts as potential pro-inflammatory modulators in IPF.[26] JAK1, STAT1 and STAT3 signalling were represented in another node. Of interest, TGF-β1, IL-6 and Oncostatin M have all been reported to engage JAK1, STAT1 and STAT3 signaling[27] and genetic modulation of *STAT3* is protective in the bleomycin-induced lung fibrosis model.[28] Moreover, therapeutic approaches targeting these mechanisms are currently being pursued in the context of IPF and other fibrotic indications.[29, 30] Finally, WGCNA analysis also identified translation as a prominent feature of the transcriptional signature of FF and is consistent with the highly ECM synthetic nature of myo/fibroblasts within FF.

The second bioinformatics approach aimed to identify key programmes associated with the generation of pathogenic ECM and was based on the core principle of WGCNA analysis that was adapted in order to use the first principal component of the expression of fibrillar collagen genes as a measure of underlying collagen production. We followed convention by naming this a collagen *eigengene*.[31] To the best of our knowledge, this is the first report describing the implementation of this approach to define pathways underlying fibrogenesis in IPF or indeed any other human fibrotic condition. This analysis identified around 300 genes with expression patterns that were either positively or negatively correlated with the collagen *eigengene*. It is worth mentioning that we found no significant overlap between these collagen-related genes and the list of genes differentially expressed in myofibroblasts in IPF compared to control lung reported in a recent bioRχiv scRNAseq study.[8] This suggest that scRNAseq and our LCM approach could be considered complementary strategies to interrogate different aspects of IPF pathomechanisms.

Interestingly, there was only one interaction network within the 300 genes comprising known protein-protein-interactions validated by multiple publications, resulting in a composite module. It is worth commenting that four of the top 15 correlated genes in the dataset were inversely correlated with the collagen *eigengene* (*ATG4B, PLEC, SELM and WIZ*) and may reflect mechanisms that are downregulated or inhibited during the course of collagen production. Of interest, *Atgb4*-deficient mice exhibit more extensive and severe fibrosis with increased accumulation of collagen relative to control mice in the bleomycin model of lung fibrosis.[32]

Cytoskeletal re-organisation, TGFβ-1 signalling, ECM production and cell-matrix interactions are fundamental properties expected of myo/fibroblasts. These mechanisms mapped strongly to the collagen *eigengene* signature and therefore provided important validation of the utility of the LCM and collagen *eigengene* approach to identify biology of relevance to the primary functions of these cells. It is worth highlighting the presence of a TGF-β1 signalling sub-module, which includes both the TGF-β1 ligand and the αvβ5 integrin, which has been implicated in the activation of latent TGF-β1 *in vitro*.[33, 34] To the best of our knowledge, this represents the first human tissue-based evidence demonstrating a link for this TGF-β1-integrin axis and fibrogenesis in IPF.[33, 35]

RhoA signalling again featured strongly within this network and was positively correlated with the collagen *eigengene*. This network also comprised two guanine nucleotide exchange factors involved in the regulation of the activation of RhoA signalling, *ARHGEF2* and *AKAP13/LBC*, which were also positively correlated with the collagen *eigengene*. Interestingly, *AKAP13/LBC* has recently been identified as a susceptibility gene in IPF.[36] While expression of AKAP13 was reported to be highest in damaged epithelium and lymphoid cells, expression was also noted in cells within FF. Moreover, ablation of AKAP13 expression in cardiac fibroblasts strongly impinges on collagen production in response to a pro-fibrotic challenge[37] so that *AKAP13/LBC* may represent both a susceptibility gene and potential drug target for inhibiting pro-fibrotic signalling.

Hand curation of this composite module also identified less predictable elements, such as fatty acid (FA) metabolism, including the gateway enzyme for FA synthesis, *FASN*, which was inversely correlated with the collagen *eigengene*. The current literature indicates both pro- and anti-fibrotic roles for altered FA metabolism in the context of pulmonary fibrosis.[38, 39] Whereas, endogenous nitrated FAs exert anti-fibrotic effects by up-regulating peroxisome proliferator-activated receptor γ (PPARγ) in human lung fibroblasts another study showed that FASN is required for pro-fibrotic TGF-β signalling.[40]

The observation that the collagen *eigengene* network also comprised a small TSC2/RHEB module, a prominent intrinsic regulator of mTORC1 activity, was of particular interest in light of the recent evidence highlighting a key role for the mTORC1/4E-BP1 axis as a core fibrogenic pathway downstream of TGF-β1.[14] The current LCM FF transcriptome and our previously published RNASeq dataset of TGF-β1-stimulated pHLFs, further revealed that *RHEB* was positively correlated with the collagen *eigengene*, whereas *TSC2* was inversely correlated, supporting a potential model of persistent mTORC1 activation. The specific mechanism by which TGF-β1 promotes the activation of mTORC1 remains poorly understood and it is also recognised that the TSC2/RHEB signalling can act independently of mTORC1.[41] Using *RHEB* CRISPR-Cas9 gene editing, we now provide strong evidence for a key role for *RHEB* in mediating both TGF-β1-induced mTORC1 activation and downstream collagen deposition in mesenchymal cells reflecting IPF and other disease settings, including the stromal reaction in cancer. It is also worth commenting that the lack of any correlations for TSC2 and *RHEB* with the collagen *eigengene* in previously published datasets based on RNA extracted from whole IPF lungs, the bleomycin model and recent scRNAseq studies of IPF lung[7, 18] emphasizes the power of the LCM approach adopted in the present study.

In summary, the findings reported in this study provide strong support for the experimental and bioinformatics approaches adopted to identify key transcriptional programmes associated with FF. As well as confirming expected pathways associated with myofibroblast biology, this analysis provides unprecedented biological insights into some of the key signalling networks associated with fibrogenesis *in situ* and place TSC2/RHEB downstream of TGF-β1 signalling as being central to the fibrogenic response in multiple mesenchymal cells. Targeting this axis may hold therapeutic potential in the context of IPF and other conditions associated with pathogenic fibrosis, including the stromal reaction in cancer.

## AUTHOR CONTRIBUTIONS contributions

D.G. performed *in vitro* experiments, interpreted data, prepared Figures and participated in drafting of the manuscript and reviewed and approved the final version

A.R.T. designed the bioinformatics analysis workflow, participated in drafting the manuscript and reviewed and approved the final version

M.P. performed *in vitro* experiments and analysed data and reviewed and approved the final manuscript

P.F.M. supervised the study and identified IPF FF and reviewed and approved the final manuscript

L.M.E. designed the bioinformatics analysis workflow and approved the final manuscript

R.H. and G.M. performed laser capture microdissection (LCM) of IPF FF and microarray analysis and approved the final manuscript

R.J.M. obtained ethical approvals, provided patient tissue, participated in the identification of IPF FF and reviewed and approved the final manuscript

A.J.F. and T.M.M. obtained ethical approvals, consented patients and provided IPF patient tissue, collected clinical data and reviewed and approved the final manuscript

R.E.H. isolated and characterized cancer associated fibroblasts (CAFs) from patients with lung adenocarcinoma (TRACERx clinical study) and approved the final manuscript

M.J-H. on behalf of the TRACERx consortium, obtained ethics approval, provided lung adenocarcinoma patient tissue, collected clinical data and approved the final manuscript

R.P.M. conceived the study and reviewed and approved the final manuscript

R.C.C. and A.D.B. conceived and designed the study, interpreted data, drafted the manuscript and reviewed and approved the final version

## ACKNOWLEDGEMENTS

The authors acknowledge several institutions for the provision of human IPF lung tissue, including the Institute of Transplantation, Freeman Hospital, Newcastle Upon Tyne (n=9 patients undergoing lung transplantation for end–stage disease); University College London Hospitals (n=3 patients undergoing surgical lung biopsy for diagnostic purposes) and the Brompton and Harefield NHS Trust (n=1 patient undergoing lung transplantation for end– stage disease). The authors thank members of the lung TRACERx consortium whose study enabled the derivation of the CAFs used in this study. The authors thank Dr Pascal Durrenberger (Centre for Inflammation and Tissue Repair, UCL Respiratory, University College London, London, WC1E 6JJ, UK) for the images provided in Figure 1a.

## SUPPLEMENTARY TABLE

### Table S1: Probe intensity and analysis.

Table of the probesets that constitute the fibroblastic focus gene signature (probesets detected in >7 patients) with the corresponding gene symbol and gene title (column B and C) as well as the log 2 intensity values for each patient (column D-P) and the WGCNA module they were associated to (column Q). The R values of the probesets correlating to the collagen eigengene (p<0.01) are presented in column R.

